# Scavenger: A pipeline for recovery of unaligned reads utilising similarity with aligned reads

**DOI:** 10.1101/345876

**Authors:** Andrian Yang, Joshua Y. S. Tang, Michael Troup, Joshua W. K. Ho

## Abstract

**Motivation:** Read alignment is an important step in RNA-seq analysis as the result of alignment forms the basis for further downstream analyses. However, recent studies have shown that published alignment tools have variable mapping sensitivity and do not necessarily align reads which should have been aligned, a problem we termed as the false-negative non-alignment problem.

**Results:** We have developed Scavenger, a pipeline for recovering unaligned reads using a novel mechanism which utilises information from aligned reads. Scavenger performs recovery of unaligned reads by re-aligning unaligned reads against a putative location derived from aligned reads with sequence similarity against unaligned reads. We show that Scavenger can successfully recover unaligned reads in both simulated and real RNA-seq datasets, including single-cell RNA-seq data. The reads recovered contain more genetic variants compared to previously aligned reads, indicating that divergence between personal and reference genomes plays a role in the false-negative non-alignment problem. We also explored the impact of read recovery on downstream analyses, in particular gene expression analysis, and showed that Scavenger is able to both recover genes which were previously non-expressed and also increase gene expression, with lowly expressed genes having the most impact from the addition of recovered reads. We also found that the majority of genes with >1 fold change in expression after recovery are categorised as pseudogenes, indicating that pseudogene expression can be affected by the false-negative non-alignment problem. Scavenger helps to solve the false-negative non-alignment problem through recovery of unaligned reads using information from previously aligned reads.

**Availability:** Scavenger is available via an open source license in https://github.com/VCCRI/Scavenger/

## 1 Introduction

Read alignment is the process of mapping high-throughput sequencing reads against a reference genome or transcriptome to identify the locations from which the reads originate. This step is typically one of the first steps in the analysis of RNA sequencing (RNA-seq) data prior to downstream analyses such as variant calling and gene expression analysis. There have been a number of published tools which have been developed to perform RNA-seq alignment, such as HISAT2 [13], STAR [7] Subread [17], CRAC [21], MapSplice2 [24] and GSNAP [25]. More recently, new alignment-free tools have been developed specifically for gene expression analysis which skips the alignment of reads to the reference and instead performs pseudoalignment. However, these alignment-free tools are only applicable to specific types of analyses and have limitations compared to traditional alignment methods [10]. The correctness of alignment programs are crucial to the accuracy of the downstream analyses. Unfortunately, previous studies have shown that while these tools have low false positive rates, they do not necessarily have low false negative rates [3, 1]. This means that while many of the reads were likely to be correctly aligned, there are still many incorrectly unaligned reads which should have been aligned. These incorrectly unaligned reads, or false negative non-alignments, adversely affect the accuracy of the alignment produced and can also affect the result of downstream analyses, such as variant calling, indel (insertion-deletion) detection and gene fusion detection [1].

There are a number of factors which contribute to the false negative non-alignment problem. One such factor is the type of algorithm utilised in the alignment tool. In order to efficiently perform alignment against a typically large reference genome in an acceptable amount of time, and to account for splicing events inherent in RNA-sequencing data, many alignment tools use heuristic-based matching of seed sequences generated from read sequences. Due to the typically short length of a seed sequence and the existence of repetitive regions within the genome, there may be multiple locations assigned to a given read which results in the alignment tool excluding the read due to ambiguity – a problem known as multi-mapping reads. Another factor which causes false negative non-alignment problems is the divergence between the reference genome and the personal genome of the organism being sequenced. The reference genome is typically constructed from a small number of samples and thus will only represent a limited degree of the organism’s diversity. Alignment of reads to the reference genome will thus be imperfect due to natural variation present in an individual organism. While alignment tools do take into account the variability between the reference genome and an individual’s genome by allowing for mismatches, insertions and deletions during alignment, they are unable to handle a substantial degree of genetic variation, such as hyper-edited sites, gene fusion and trans-splicing.

Correcting for a false negative non-alignment problem is much more difficult compared to correcting false positive reads. For false positive reads, there are a number of strategies which can be employed to help filter these type of reads, such as by removing lower quality alignments, removing reads with multiple alignment locations and re-aligning reads with a more specific alignment tool. Recovering false negative reads, on the other hand, is not as straightforward as it is not possible to identify their putative alignment region in the genome. One possible strategy for solving the false negative non-alignment problem is to tune the parameters used for alignment in order to maximise the amount of reads aligned, such as by increasing the threshold for multi-mapping reads and/or increasing the number of mismatches allowed. However, this approach is limited as there is no ground truth in real data to help with optimisation, and increasing the number of reads aligned will also result in an increase in the number of false positive reads. Another strategy for solving the false-negative non-alignment problem is by incorporating variation information during alignment, in the form of utilising alternate loci sequences within the reference genome [15] or integration of a single nucleotide polymorphism database to the reference [13], to help minimise the effect of divergence of the personal genome compared to the reference genome. This approach is also limited as it requires existing variation information, which may not be available in non-model organisms.

We have recently applied the idea of Metamorphic Testing – a software testing technique designed for the situation where there is an absence of an oracle (a method to verify the correctness of any input) – for performing software testing on the STAR sequence aligner [23]. Metamorphic testing involves multiple executions of the program to be tested with differing inputs, constructed based on a set of relationships (Metamorphic relations – MR), and checking that the outputs produced satisfy the relationships [5, 6]. In our previous study [23], we developed an MR to test the re-alignability of previously aligned reads in the presence of irrelevant ‘control’ chromosomes constructed from previously unaligned reads. We discovered that a non-trivial amount of reads that were previously aligned to the reference genome were now aligned to the control chromosomes consisting of reads which were unable to be aligned to the reference. Further investigation indicated that some of the unaligned reads have high similarity to the aligned reads, indicating the possibility of these reads being false negative non-alignments.

In this chapter, we aim to tackle the problem of false-negative non-alignments by taking inspiration from our previous work on metamorphic testing. We have developed Scavenger, a pipeline designed to recover incorrectly unaligned reads by exploiting information from reads which are successfully aligned. We applied the Scavenger pipeline on a number of simulated and actual RNA-seq datasets, including both bulk (normal) and single-cell RNA-seq datasets, and demonstrated the ability of Scavenger in recovering unaligned reads from these datasets. We then analysed the impact of adding these recovered reads on downstream analyses, in particular gene expression analysis, and discovered that lowly expressed genes, in particular genes of the pseudogenes category, are more affected by the false-negative non-alignment problem. We also verified that the divergence between the personal genome and the reference genome is a contributing factor to the false-negative non-alignment problem and showed that Scavenger is able to recover reads which are unaligned due to higher degree of variability within the reads sequence.

## 2 Methods

Scavenger is a python-based pipeline designed to recover unaligned reads by utilising information from aligned reads. The pipeline takes in sequencing reads in FASTQ format as the input, along with a reference genome sequence in FASTA format and a corresponding index for the alignment tool built using the reference genome. There are 4 main steps in the Scavenger pipeline – source execution of alignment tool, follow-up execution using aligned reads as input and unaligned reads as index, consensus filtering of follow-up execution result to obtain putative alignment location, and re-alignment of unaligned reads to the reference genome (Figure 1). The unaligned reads which are able to be successfully re-aligned back to the genome are then re-written back to the alignment result from the source execution.

**Figure 1:**
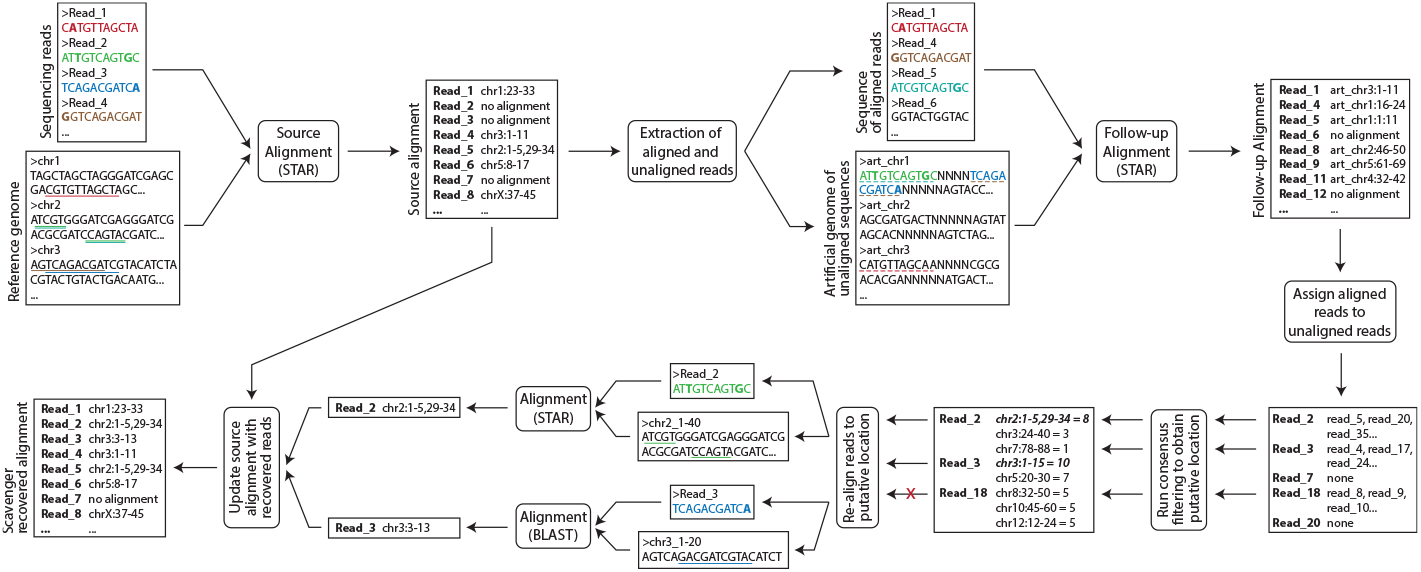
Overview of the Scavenger pipeline. Scavenger first aligns sequencing reads against the reference genome using the STAR alignment tool in the source execution step. Scavenger then extracts both the aligned and unaligned reads from the source alignment result and creates a sequencing reads file based on the aligned reads and an artificial genome file containing chromosomes built using sequences of unaligned reads. The sequences of aligned reads are then aligned to the artificial genomes using the same alignment tool from the source execution (STAR) in the follow-up execution steps to find aligned reads which have similar sequences to unaligned reads. Next, consensus filtering is performed to select putative sites for re-aligment based on where the majority of aligned reads originate from in the reference genome. Finally, re-alignment is performed for unaligned reads which pass consensus filtering and the source alignment result is updated based on the result of re-alignment.

### 2.1 Source execution

The first step of the Scavenger pipeline is the source execution where sequencing reads are aligned to the reference genome using a sequence alignment program. The alignment program used must satisfy the three properties which are required to validate the metamorphic relation underlying the read recovery pipeline – deterministic alignment, realignability of mapped reads, and non-realignability of unmapped reads. Currently, STAR is utilised for aligning RNA sequencing reads in the Scavenger pipeline as it has been previously evaluated as being a reliable general-purpose RNA-seq aligner, with good default performance, [3] as well as satisfying the three properties above [23]. The source execution step can be skipped if the user has previously performed alignment of sequencing reads by passing in the alignment file produced in either SAM or BAM format as input to the Scavenger pipeline.

### 2.2 Follow-up execution

In the follow-up execution step, both aligned and unaligned reads are first extracted from the alignment file produced during source execution. For reads which have been successfully and uniquely aligned, a sequencing reads file (in FASTQ format) is created using the reads’ sequence and qualities retrieved from the alignment records. In the case of reads which did not align to the reference genome, reads with identical sequences are first grouped together in order to minimise computational complexity and to reduce the potential location for alignment. The unique unaligned sequences are then extended with spacer sequences (sequence of N nucleotides) in order to form sequence bins of equal length and to ensure that aligned reads do not align between two unaligned sequences. These sequence bins are concatenated into artificial chromosomes and stored into a new temporary genome file. Depending on the alignment program utilised, a new index will then need to be created based on the temporary genome containing the artificial chromosomes prior to alignment. Finally, sequencing reads of previously aligned reads are aligned to the temporary genome containing unaligned read sequences using the alignment tool used in source execution. In the current Scavenger pipeline, STAR is again utilised in the follow-up execution with a number of extra parameters in order to disable spliced alignment to ensure that input reads only align to one unaligned read sequence and to remove the restriction of the number of locations (i.e. unaligned read sequence) that the input reads can align to in the temporary genome.

### 2.3 Consensus Filtering

The next step of the Scavenger pipeline is consensus filtering. Reads which align during the follow-up execution step are extracted from the alignment file produced from the previous step to obtain information regarding similarity between reads aligned during source execution and reads which did not align during source execution. Each unaligned sequence may have alignments to multiple aligned reads from the source execution. As these aligned reads may be aligned to different regions in the reference genome, consensus filtering is performed to select putative sites for re-alignment. For each unaligned sequence, intervals are created based on the reference genome location of previously aligned reads that align to the unaligned sequence. Overlapping intervals are then merged to form longer intervals to both reduce the number of putative sites and to increase the support for the interval to be selected as a putative site. An interval is considered as being a putative site if there is more than one read within the interval and the level of support for the interval (i.e. the number of aligned reads that fall within the interval) is greater than the consensus threshold, which is set to 60% of the number of previously aligned reads that align to the unaligned sequence by default. During this step, there is also an optional filtering criteria that can be utilised to remove unaligned sequences which likely originate from a low complexity region or tandem repeat region. The filtering method is based on the tandem repeat detection step used in the ROP tool [19], which uses MegaBLAST [4] to align reads against a repeat sequence database, such as RepBase [2].

### 2.4 Re-alignment

The final step is the re-alignment step where unaligned sequences which pass the filtering steps are re-aligned to the reference genome using the putative location obtained from reads aligned during source execution as a guide. For each unaligned sequence, the reference genome sequence around the putative location (extended 100 base pairs at both the start and the end of the putative location) is extracted and stored as the new genome for aligning the unaligned sequence. Alignment of the unaligned sequence is then performed against the new genome using either MegaBLAST or STAR depending on the putative location of the unaligned sequence originate from the unspliced alignment or from the spliced alignment during the source execution, respectively. MegaBLAST is utilised for unspliced alignment due to its high sensitivity, though a strict parameter of 64% overlap and 85% query identity (which replicates the result of STAR alignment) is also utilised to reduce the false positive recovery of sequences. Unaligned sequences which are successfully and uniquely aligned back to the reference genome are then added back to the alignment file of the source execution by modifying the alignment records of previously unaligned reads whose sequence matches the recovered unaligned sequence.

### 2.5 Parallelising Scavenger

Both the consensus and re-alignment steps of the Scavenger pipeline are computationally expensive due to the potentially large number of unaligned reads to be processed. However, the processing of the inputs are independent to each other thus allowing for parallelisation of processing unaligned reads in order to reduce the overall runtime of the pipeline. Scavenger takes advantage of Python’s built-in multiprocessing library in order to parallelise the consensus and re-alignment steps across the available CPU cores of the machine.

To enhance the scalability of Scavenger, a framework has been provided to enable parallel computation of a read recovery session on cloud computing resources. Cloud computing enables convenient, on-demand network access to a shared pool of configurable computing resources [20]. Central to the model of cloud computing is the virtualisation of computing resources to enable sharing of pooled resources. These resources can be commissioned and decommissioned as the user requires. Scavenger has a framework that employs the resources offered by the cloud provider Amazon Web Services (AWS). The cloud provider enables the user, using their own account credentials, to create a number of computing ‘‘instances”, which are the virtual machines upon which the user can perform their computational workload. In the case of AWS, such resources are termed “EC2 instances”. An instance typically can be provisioned within minutes of the user request, and the user is charged by the hour. Some cloud providers, such as AWS, offer reduced price “spot” instances at a greatly reduced price, such that the user places a “bid” for a spot instance on the proviso that the instance will be terminated should the current market price for the instance exceed the initial bid price. To minimise the cost for users, Scavenger utilises AWS spot instances.

The cloud computing feature of Scavenger, after initial configuration on the user’s controlling computing resource, uses the AWS EC2 cloud instances to perform the various steps of read recovery, and also uses AWS cloud storage (S3) to store test data and results. The Scavenger cloud processing feature co-ordinates all interactions with the cloud resources, with logging information stored both locally and on the cloud. The user can elect to have a large job to be spread among a number of cloud instances, with Scavenger creating the instances and distributing the work load evenly amongst the instances. The cloud computing feature of Scavenger is optional, and the user can elect to use their own computing resources if desired.

### 2.6 Datasets

Three different types of RNA-seq datasets – simulated, normal (bulk) and single-cell – were utilised to evaluate the Scavenger pipeline. The simulated datasets were obtained from a previous study [3] which generated 3 sets of simulated RNA-seq datasets from the hg19 reference genome using BEERS simulator [8] with varying parameters to emulate different level of dataset complexity. As the simulated datasets were formatted in FASTA format, high quality scores were added to each of the simulated reads to produce corresponding FASTQ files. These files were then input into Scavenger for both source alignment and read recovery with either STAR v2.5.3a or Subread v1.6.0 as the alignment tool. The GRCh37.p13 reference genome was obtained from GENCODE [9] and modified to contain reference chromosomes only, and used to create the indexes for each alignment tool. For STAR specifically, the annotation file was extracted from a previous study [3] and utilised in index creation to help increase the accuracy for alignment across splice junctions. In the evaluation of alignment results for simulated datasets, we used the analysis script that was used in the previous study [3] to analyse the correctness of the alignment results.

The normal and single-cell RNA-seq datasets were obtained from publicly available human and mouse datasets which were deposited to the NCBI Sequence Read Archive [16] (Table 1). Pre-processing of the datasets was performed using Trimmomatic v0.36 to remove low quality sequence and short reads. The pre-processed datasets were then analysed by Scavenger using STAR v2.5.3a as the alignment tool in the source execution and for re-alignment of spliced reads, together with BLAST v2.6.0 for re-alignment of unspliced reads. Indexes used for aligning of both human and mouse datasets were generated from GRCh38 and GRCm38 reference genomes respectively, which were obtained from GENCODE together with the corresponding annotation files (version 27 for human and version 15 for mouse). As before, annotation was used to augment the index to increase accuracy for alignment. The Repbase database [2] was also utilised to remove low complexity reads and reads from repetitive regions. For human datasets, the simple, humrep and humsub sequence files from Repbase were concatenated and used to create a BLAST database. Reads that passed consensus were aligned to this database and the aligned reads that have a minimum of 90% sequence identity and 80% sequence coverage were removed for further processing in Scavenger. A similar approach was used for the mouse datasets, but the simple and mousub sequence files were used instead. For mouse strain analysis, strain-specific VCF files for non-reference mouse strains con-taining SNPs derived against the reference C57BL/6J mouse genome were downloaded from the Mouse Genome Project (MGP) [12].

**Table 1:** List of datasets used for Scavenger testing and evaluation. The datasets are divided into three sections: 1. Datasets from selected non-reference mouse strain, 2. Normal (bulk) RNA-seq dataset from either human or mouse, and 3. Single-cell RNA-seq dataset from mouse.

| Accession ID | Samples ID | Organism | Tissue/Source |
| --- | --- | --- | --- |
| SRP039411 | SRR1182782 - SRR1182783 | Mus Musculus | Liver |
| ERP000614 | ERR032989 - ERR032991; ERR032997 - ERR032998;<br>ERR033006 - ERR033009; ERR033017 - ERR033019 | Mus Musculus | Brain |
| SRP020636 | SRR826292 - SRR826299; SRR826308 - SRR826315;<br>SRR826340 - SRR826347; SRR826356 - SRR826363 | Mus Musculus | Liver |
| SRP068123 | SRR3087147 - SRR3087158; SRR3087171 -<br>SRR3087176 | Mus Musculus | Hippocampus |
| SRP013610 | SRR504764 - SRR504766 | Mus Musculus | Eye |
| SRP076218 | SRR3641982 - SRR3641983; SRR3641990;<br>SRR3642003 - SRR3642005; SRR3642012 -<br>SRR3642014 | Mus Musculus | Heart |
| SRP045630 | SRR1554415 - SRR1554417 | Mus Musculus | Retina |
| SRP016501 | SRR594393 - SRR594401 | Mus Musculus | Brain; Colon; Heart;<br>Kidney; Liver; Lung;<br>Skeletal Muscle; Spleen;<br>Testes |
| SRP075605 | SRR3578721 - SRR3578725 | Homo sapiens | Fibroblasts |
| SRP122535 | SRR6337339 - SRR6337344 | Homo sapiens | ESC |
| SRP013027 | SRR4422503 - SRR4422506; SRR4422535 -<br>SRR4422538; SRR4422626 - SRR4422629 | Mus Musculus | Hindbrain; Limb; Heart |
| SRP045452 | 80 randomly selected samples | Mus Musculus | Hippocampus |

For running the alignment using STAR, the following command is used: `STAR --runThreadN <threads> <aligner_extra_args> --genomeDir <genome_index> --readFilesIn <read_files> --outFileNamePrefix <output_prefix>`. As for running the alignment using subread, the following command is used: subread-align -T `<threads> -t 0 <aligner_extra_args> -i <genome_index> <read_files> -o <output_file> <bam_option>`. And lastly, for running the alignment using BLAST, the follow command is used: `blastn -query unmapped_read -subject target_genome -task megablast -perc_identity <identity> -qcov_hsp_perc <coverage> -outfmt “17 SQ SR” -out <sam_output> -parse_deflines`. During follow-up alignment using STAR, the following parameters are additionally used: `--outFilterMultimapNmax <num_reads> --alignIntronMax 1 --seedSearchStartLmax 30`.

## 3 Results

### 3.1 Recovery of reads on simulated data

To evaluate the ability of the Scavenger pipeline to recover false-negative non-aligned reads, we first tested Scavenger using previously published human simulated data. The varying level of complexity of the simulated datasets represents the degree of divergence between the sequencing reads generated compared to the reference genome, ranging from low polymorphism and error rate (T1), moderate polymorphism and error rate (T2) and high polymorphism and error rate (T3). The results of the source execution of STAR with default parameters are consistent with the previously published result, with >99% of reads being aligned in both T1 and T2 and >90% of reads being aligned in T3 (Table 2). After running the Scavenger pipeline, we were able to recover between 4-30% of the previously unaligned reads in the three datasets, resulting in an increase of aligned reads ranging from ~1,500 to ~160,000. The majority of reads recovered by Scavenger are aligned in the correct position, with 79.4% of reads being correctly recovered in T1 and >98% of reads being correctly recovered in T2 and T3.

**Table 2:** Alignment statistics for simulated datasets before and after recovery of reads with Scavenger using default parameters for STAR. The result shown is an average from 3 samples.

| Dataset | Source execution |  |  | Scavenger pipeline |  |  | Unaligned reads recovered | % recovered reads correct | % recovered reads incorrect |
| --- | --- | --- | --- | --- | --- | --- | --- | --- | --- |
|  | Aligned correctly | Aligned incorrectly | Unaligned | Aligned correctly | Aligned incorrectly | Unaligned |  |  |  |
| T1 | 9,671,586 | 8,022 | 33,486 | 9,672,770 | 8,330 | 31,994 | 1,492 | 79.4 | 20.6 |
| T2 | 9,617,585 | 17,163 | 56,827 | 9,634,469 | 17,496 | 39,610 | 17,217 | 98.1 | 1.9 |
| T3 | 8,595,549 | 67,559 | 933,274 | 8,753,899 | 67,995 | 774,488 | 158,786 | 99.7 | 0.3 |

The difference in the number of aligned reads between the three datasets can be explained by the degree of divergence between the sequencing reads and the reference genome; and the limitation of the alignment tool in aligning reads which display a high degree of polymorphism. The simulated sequencing reads in both T1 and T2 have high homology to the reference genome due to the lower degree of polymorphism and error rate introduced meaning that the majority of these reads will be accurately mapped to the reference genome with a very small number of mismatches during alignment. In contrast, the sequencing reads in T3 – with the higher polymorphism and error rate – have a much higher degree of divergence compared to the reference genome thus resulting in more mismatches during alignment and therefore causing it to fail to be aligned. The Scavenger pipeline is able to recover more reads in T2 and T3 compared to T1 due to the greater number of aligned reads that contain mutations within the sequence. During follow up execution, Scavenger exploits the fact that these aligned reads will have closer similarity to the unaligned reads, which will also contains mutations, therefore resulting in the alignment of the aligned reads to the unaligned reads to obtain the putative location for the unaligned reads for recovery.

Another method to solve the false-negative non-alignment problem is to adjust the parameters of the alignment tool utilised in order to allow alignment of reads with a higher degree of polymorphism. As has been shown previously, alignment of the simulated datasets using STAR with optimised parameters results in >99.2% of the reads being aligned, with T1 and T2 reaching nearly 99.9% of reads being aligned (Table 3). The Scavenger pipeline is unable to obtain the high degree of alignment achieved with parameter optimisation due to limitations in Scavenger’s approach to recover reads. Since Scavenger utilises information from aligned reads to find the putative location of unaligned reads for recovery, it is not possible to recover any unaligned reads from regions which have no read alignments. As such, the reads that the Scavenger pipeline is able to recover are reads from regions which already have alignment. This is unlike parameter optimisation, which allows for alignment with a higher threshold of mismatches in any region irrespective of whether there was alignment in the region. This observation can be seen in the high degree of overlap (>96.5%) of the reads recovered by the Scavenger pipeline compared to the reads recovered by optimised parameters. The Scavenger pipeline is still able to recover some reads which are unaligned with optimised parameters, particularly in T3 where Scavenger recovered ~9.75% of previously unaligned reads. Unlike Scavenger recovery with default parameters, the majority of recovered reads after alignment with optimised parameters are incorrectly aligned in both the T1 and T2 datasets. Given the very high degree of alignment in these lower complexity datasets, it is likely that the unaligned reads are reads which can align to many locations in the genome and thus correctly recovering these reads is very difficult and error prone. These results indicate that parameter optimisation provides a solution to the false-negative nonalignment problem, performing better than Scavenger. However, given that performing parameter optimisation is not trivial due to lack of ground truth in real datasets, these results also show that Scavenger can be utilised as an alternative to help recover false-negative non-aligned reads.

**Table 3:** Alignment statistics for simulated datasets before and after recovery of reads with Scavenger using optimised parameters for STAR. The result shown is an average from 3 samples.

| Dataset | Source execution |  |  | Scavenger pipeline |  |  | Unaligned reads recovered | % recovered reads correct | % recovered reads incorrect |
| --- | --- | --- | --- | --- | --- | --- | --- | --- | --- |
|  | Aligned correctly | Aligned incorrectly | Unaligned | Aligned correctly | Aligned incorrectly | Unaligned |  |  |  |
| T1 | 9,673,309 | 6,861 | 15,660 | 9,673,362 | 6,948 | 15,519 | 141 | 37.8 | 62.2 |
| T2 | 9,643,573 | 14,570 | 11,237 | 9,643,715 | 14,675 | 10,990 | 246 | 55.5 | 44.5 |
| T3 | 9,437,748 | 75,395 | 83,687 | 9,445,855 | 75,448 | 75,527 | 8,160 | 99.4 | 0.6 |

We also performed a comparison of the Scavenger pipeline against a recently published tool, Read Origin Protocol (ROP), which is primarily designed to identify the origin of unaligned reads [19]. The ROP tool consists of 6 steps, with each step designed to identify different causes for unaligned reads: reads with low quality, lost human reads, reads from repeat sequences, non-colinear RNA reads, reads from V(D)J recombination and reads belonging to microbial communities. The result of running ROP on the simulated dataset shows that ROP is able to identify an average of ~29,000 reads in the T1 and T2 datasets, and ~58,500 reads in T3 dataset (Table 4). In particular, the majority of reads in the T1 and T2 dataset are correctly identified as lost human reads, while the majority of reads in T3 dataset are incorrectly identified as immune reads. Checking the correctness of ROP identified reads is not straightforward given that most steps within ROP does not produce alignment information. Thus, correctness testing was performed only on the genome-based alignment information produced during the lost reads steps. The result of the correctness testing shows that >92.6% of the reads identified by ROP are incorrectly aligned (Table 5).

**Table 4:** Unaligned reads identified by ROP in the simulated dataset. The result shown is an average from 3 samples.

| Dataset | Unaligned<br>reads<br>identi-<br>fied | Low<br>Quality<br>Reads | Low<br>Com-<br>plexity<br>Reads | rRNA<br>reads | Lost<br>Reads | Repeat<br>reads | NCL<br>Reads | Immune<br>Reads | Microbial<br>Reads |
| --- | --- | --- | --- | --- | --- | --- | --- | --- | --- |
| T1 | 31,469 | 1 | 188 | 251 | 30,398 | 502 | 9 | 120 | 1 |
| T2 | 27,328 | 0 | 306 | 148 | 23,508 | 1,690 | 28 | 1,639 | 9 |
| T3 | 58,544 | 3 | 2,469 | 13 | 3,085 | 7,123 | 132 | 45,696 | 24 |

**Table 5:**
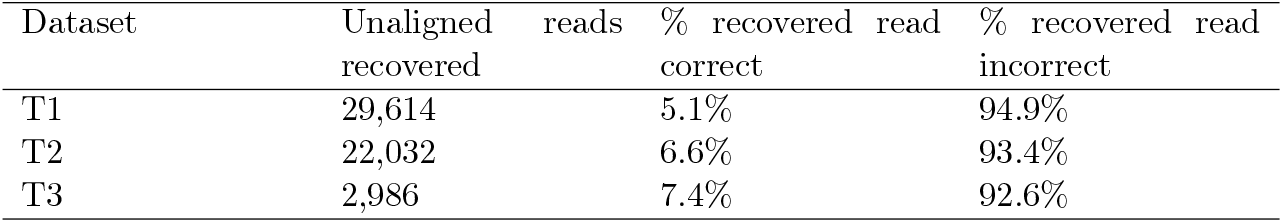
Alignment statistic for unaligned reads recovered by ROP lost read identification step. The result shown is an average from 3 samples.

### 3.2 Divergence of personal genome results in false-negative non-aligned reads

One factor which may affect the false-negative non-alignment problem is the divergence of sequences between the reference genome and personal genome which results in alignment tools being unable to properly align the reads due to the higher number of mismatches. To evaluate the ability of Scavenger in recovering these false-negative non-aligned reads which arise due to divergence of the personal genome, an experiment was devised where reads from non-reference inbred laboratory mouse strains were aligned to the reference C57BL/6J mouse genome to imitate alignment of reads from the personal genome against the reference genome. Multiple non-reference mouse strains – 129S1/SvImJ, A/J, CAST/EiJ, DBA/2J and NOD/ShiLtJ – were utilised as the genomes of these strains have previously been characterised by the Mouse Genome Project (MGP), with variations from each strain identified relative to the reference mouse genome. We collected 80 publicly available RNA-seq samples from the selected mouse strains, with each strain having a minimum of 13 samples from at least 3 different projects with varying characteristics, and performed alignment of these samples against the reference genome using STAR with default parameters. The result of the source alignments shows that there is generally a high degree of mappability of the reads, ranging from 82.2% up to 98.1%. After recovery with Scavenger, we were able to re-align ~4.75% of unaligned reads in the source execution, corresponding to an increase in the number of aligned reads ranging from 17,0 to 396,000 reads (Table 6).

**Table 6:** Alignment statistics for all RNA-seq datasets in source alignment with STAR and after recovery of reads with Scavenger. The result shown is an average of all samples per accession ID.

| Accession ID | Read length | Total reads | Source aligned reads | Source un-aligned reads | Source mappability | Scavenger aligned reads | Scavenger un-aligned reads | Scavenger mappability | Scavenger recovered reads | Unaligned reads recovered (%) |
| --- | --- | --- | --- | --- | --- | --- | --- | --- | --- | --- |
| SRP039411 | 97 | 47,077,051 | 44,052,994 | 3,024,056 | 93.6% | 44,162,052 | 2,915,000 | 93.8% | 109,057 | 3.61% |
| ERP000614 | 73 | 30,406,321 | 29,529,186 | 877,136 | 97.1% | 29,571,416 | 834,905 | 97.3% | 42,230 | 4.72% |
| SRP020636 | 93 | 10,695,056 | 10,023,946 | 671,110 | 93.8% | 10,053,119 | 641,937 | 94% | 29,173 | 4.43% |
| SRP068123 | 89 | 36,237,495 | 29,132,806 | 7,104,689 | 82.2% | 29,342,165 | 6,895,330 | 82.7% | 209,360 | 2.9% |
| SRP013610 | 54 | 21,039,752 | 20,514,308 | 525,444 | 97.5% | 20,531,454 | 508,298 | 97.6% | 17,146 | 3.19% |
| SRP076218 | 86 | 20,183,248 | 19,802,286 | 380,962 | 98.1% | 19,822,443 | 360,805 | 98.2% | 20,157 | 5.49% |
| SRP045630 | 99 | 15,931,928 | 15,550,706 | 381,221 | 97.6% | 15,578,309 | 353,618 | 97.8% | 27,603 | 7.24% |
| SRP016501 | 48 | 85,677,826 | 82,218,772 | 3,459,055 | 96.2% | 82,614,984 | 3,062,842 | 96.6% | 396,213 | 8.86% |
| SRP075605 | 51 | 30,851,404 | 29,278,793 | 1,572,611 | 95% | 29,356,220 | 1,495,184 | 95.2% | 77,427 | 5.26% |
| SRP122535 | 50 | 15,658,933 | 15,121,371 | 537,562 | 96.6% | 15,133,718 | 525,214 | 96.7% | 12,347 | 2.58% |
| SRP013027 | 100 | 28,031,517 | 26,043,731 | 1,987,786 | 92.9% | 26,092,526 | 1,938,992 | 93.1% | 48,794 | 2.49% |
| SRP045452 | 51 | 2,286,199 | 1,307,716 | 978,483 | 57.3% | 1,313,084 | 973,116 | 57.5% | 5,368 | 0.621% |

Further analysis was performed to evaluate the hypothesis that reads recovered by Scavenger have a higher degree of polymorphism due to the divergence between the ‘personal’ non-reference mouse strain genome against the reference genome. We randomly selected 1,000 unspliced reads which are aligned in the source execution and 1,000 unspliced reads recovered by Scavenger from each sample, and then calculated the number of single nucleotide polymorphisms (SNP) found within the location of the aligned reads from the list of strain-specific SNPs published by MGP against the reference mouse genome. The same analysis was then repeated a further 9 times, for a total of 10 iterations, to allow for significance testing. The majority of the reads which are either successfully aligned or recovered did not contain any of known SNPs. However, the number of reads which contain SNPs is significantly higher (p-value < 10^-27^) in the reads recovered by Scavenger compared to the reads aligned in the source execution for 4 of the 5 strains analysed (Figure 2A). Furthermore, the number of reads with a high number of SNPs (> 5) are also significantly higher (p-value < 10^-21^) in the reads recovered by Scavenger for all of the strains analysed indicating that Scavenger is able to recover reads which are more polymorphic compared to the reads aligned during the source execution (Figure 2B). These results validate the hypothesis that reads recovered by Scavenger have a higher degree of polymorphism as a result of the divergence between the personal genome and the reference genome and further demonstrates the ability of Scavenger in dealing with the false-negative non-alignment problem.

**Figure 2:**
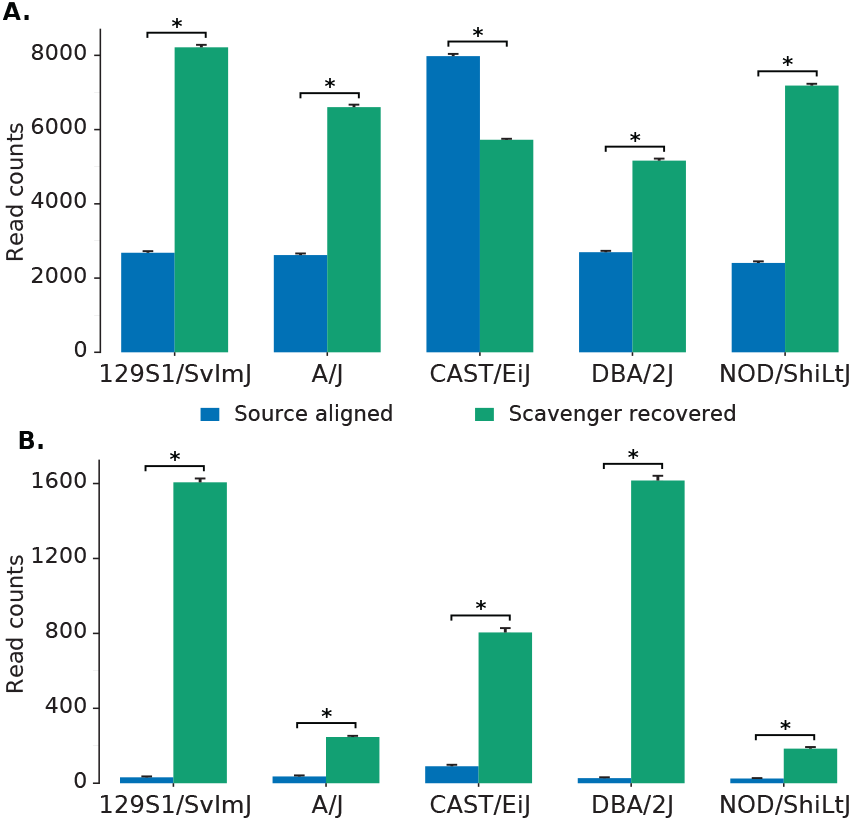
The number of reads containing SNPs found within reads aligned in source execution and reads recovered by Scavenger. A. The number of reads with ≥ 1 SNPs found within reads. B. The number of reads with high number of SNPs (> 5) found within reads.

### 3.3 Effect of Scavenger recovery pipeline on downstream analysis

While alignment of reads is an important step in RNA-seq analysis, further downstream analyses are required in order to interpret the data into meaningful results. As one of the most common applications of RNA-seq analysis is gene expression analysis, we focused on identifying the effect of adding reads recovered by Scavenger on the expression of genes. The dataset utilised for testing consisted of 23 publicly available RNA-seq samples selected from 3 separate projects of varying characteristics, with 11 samples originating from two human projects and 12 samples originating from a single mouse project. The result of source execution using STAR with default parameters shows a high degree of mappability in all datasets, ranging from ~95.9% in human datasets and ~92.9% in the mouse dataset (Table 6). After recovery of reads with Scavenger, we were able to recover ~3.1% of unaligned reads on average across the three datasets, corresponding to an increase ranging from 7,000 reads up to 102,000 reads. While the number of reads recovered are quite low relative to the number of previously aligned reads, the addition of tens and hundred of thousands of reads is still likely to affect the expression of the genes.

Gene quantification of aligned reads is performed using featureCounts [18] to produce read counts per gene, which is then normalised to reads per million (RPM). In the source alignment, the number of genes expressed, defined as having non-zero read counts, in the human datasets average to 26,000 genes, while the number of genes expressed in the mouse dataset is 25,800 genes. In Scavenger recovered alignment, we see an increase of up to 3 expressed genes per sample, indicating the ability of Scavenger to recover genes which are falsely considered as non-expressed in the source alignment (Figure 5A). The recovery of reads in previously non-expressed genes is likely due to the extension of putative alignment locations, which may introduce regions which have no alignment in the source execution. Further investigation into the reads recovered by Scavenger shows that the reads are not distributed evenly across all the expressed genes – only ~2150 and ~5900 genes receiving an increase in read counts in human and mouse datasets, respectively. The majority of genes with increased read counts do not see much change in gene expression, with only ~14 genes having more than 1 fold-change difference between source expression and recovered expression. Interestingly, genes which have substantial difference after recovery are generally genes with low expression in the source execution (log2(RPM) < 5), potentially indicating that some lowly expressed genes may actually have higher true expression than what is reported due to the alignment tool being unable to pick up these reads (Figure 4). This also has implications in further downstream analyses as lowly expressed genes are typically excluded from analysis, when instead it should not have been excluded as their true expression is actually higher.

**Figure 3:**
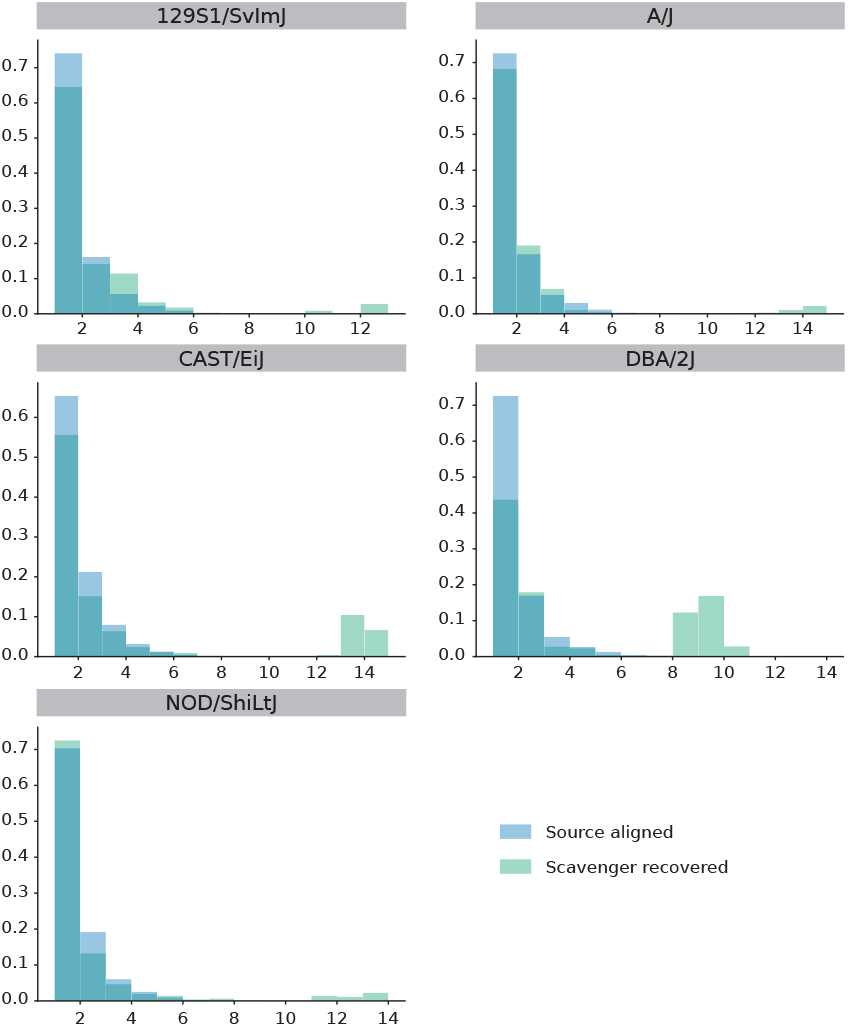
Distribution of number of SNPs found within reads recovered during source alignment and reads recovered by Scavenger. Scavenger is able to recover reads with a higher number of SNPs (>5) compared to source alignment.

**Figure 4:**
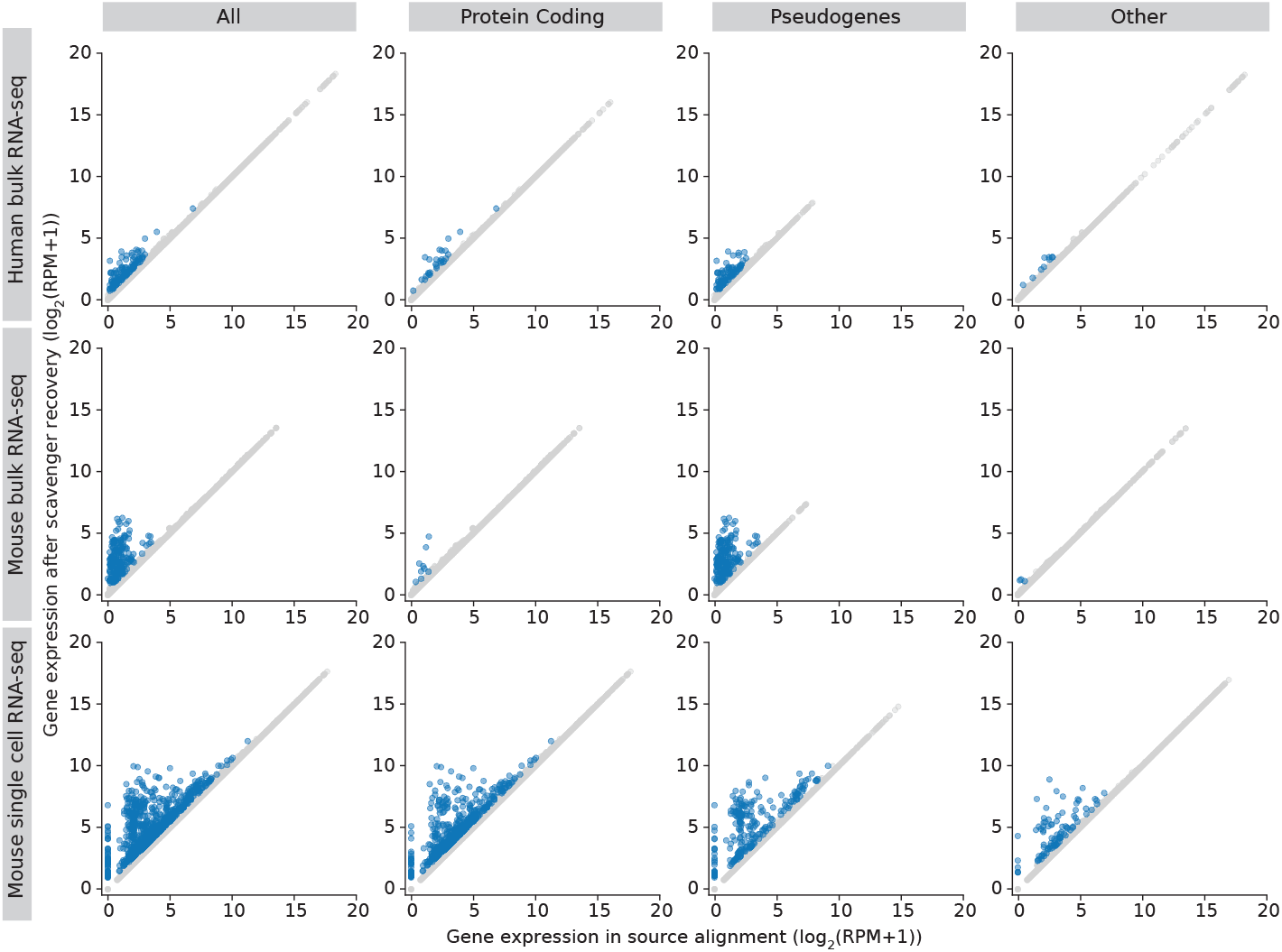
Gene expression in source alignment and after Scavenger recovery for genes whose reads are recovered. Coloured points indicates genes with expression difference of greater than 1 fold change.

We then performed further investigations into the genes with more than 1 fold-change difference after recovery to study the types of genes affected by the false-negative non-alignment problem. The majority of genes with recovered expression in the human and mouse dataset are classified as pseudogenes (>60%), with the second most frequent type being protein coding genes (22% and 9% for human and mouse dataset, respectively) (Figure 5B). Moreover, most genes with very low expression in the source alignment (log2(RPM+1) < 5) are in the pseudogenes category implying that many pseudogenes expression are likely to be under-reported due to reads originating from pseudogenes not being picked up by the alignment tool (Figure 4). Frequency analysis of the recovered genes also shows that some genes are consistently recovered across at least half of the samples in human and mouse datasets respectively, potentially indicating that these genes are harder to be picked up by the alignment tool due to its sequence being highly polymorphic. The finding that expression of pseudogenes are particularly affected by the false-negative non-alignment problem is significant as recent studies have shown that pseudogenes are incorrectly assumed to be non-functioning and actually have a role in regulating biological processes, particularly in diseases such as cancer [11, 22]. The reason that pseudogenes are more affected by Scavenger recovery is likely due to a number of factors, including the large number of mutations accumulated which results in divergence between pseudogene sequences and personal genomes; and the typically low expression of pseudogenes and correspondingly, the number of reads from pseudogenes, which therefore are more affected by increase an in reads as a result of recovery by Scavenger (Figure 7).

**Figure 5:**
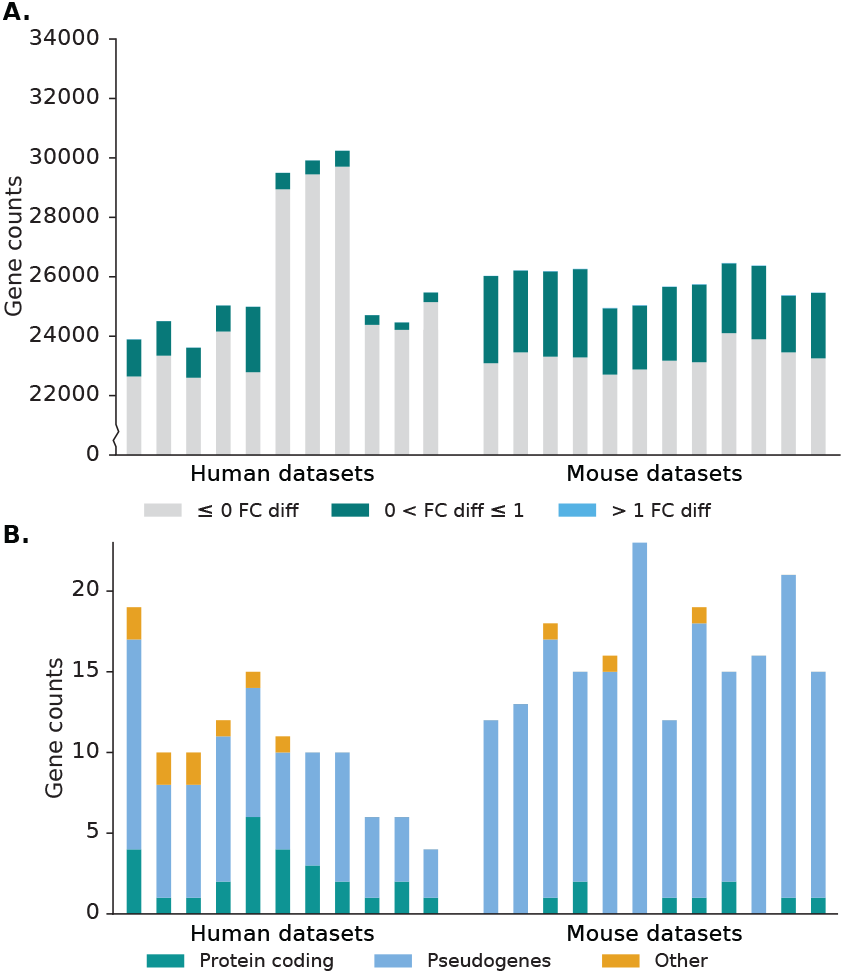
Effect of Scavenger read recovery on gene expression for normal (bulk) RNA-seq datasets. A. The number of genes whose reads are recovered by Scavenger, categorised based on the fold change in normalised expression (RPM) between source alignment and after Scavenger recovery. B. The number of genes with more than 1 fold change in normalised expression categorised based on their gene types.

**Figure 6:**
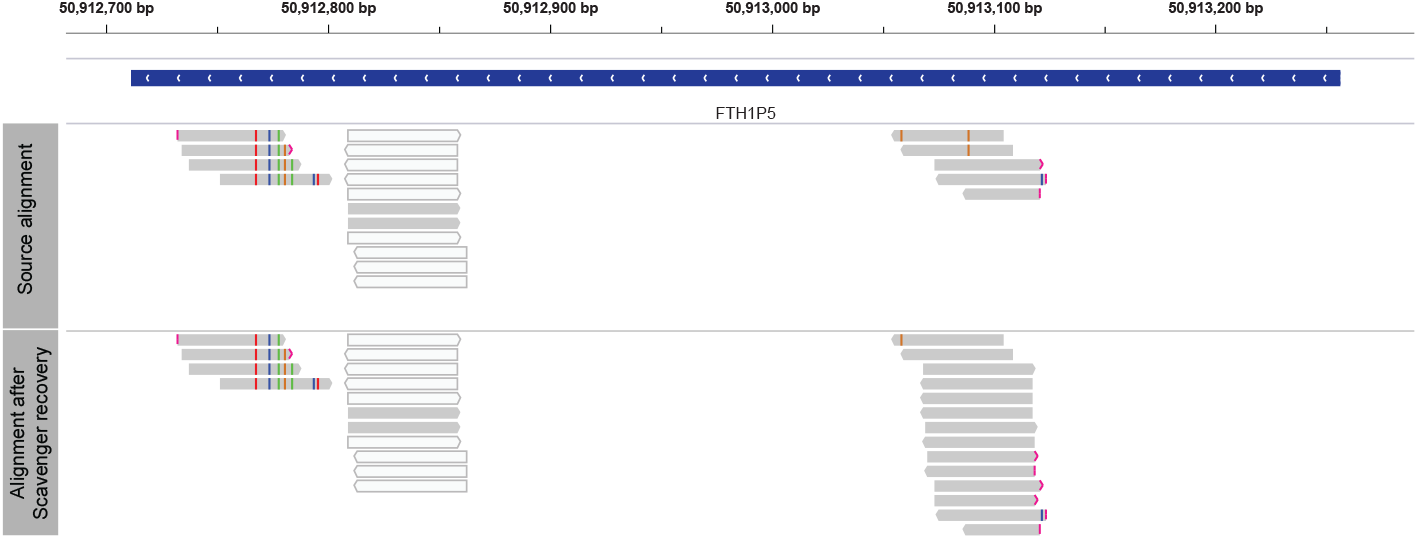
Genome browser view of alignment in source execution and after recovery with Scavenger for FTH1P5 pseudogene from one human bulk RNA-seq dataset (SRR6337341). FTH1P5 gene was chosen as it is the only pseudogene consistently recovered in at least half of the human bulk RNA-seq dataset.

**Figure 7:**
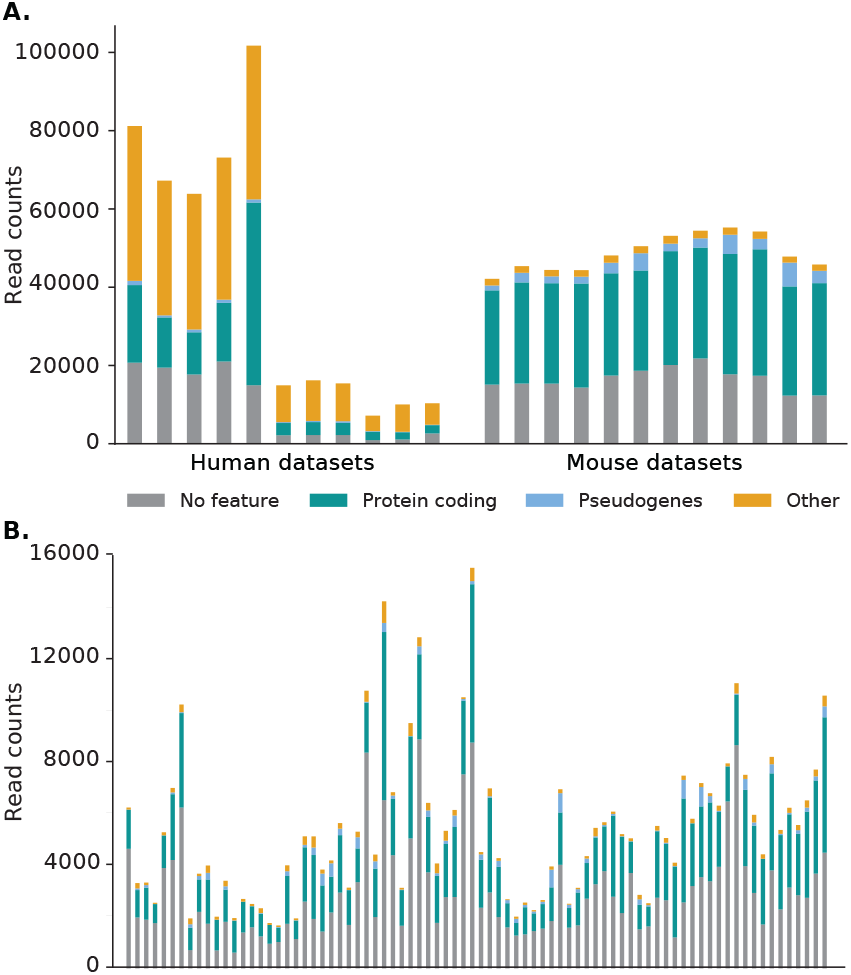
The number of reads recovered by Scavenger categorised by gene type for A. normal (bulk) RNA-seq datasets and B. single-cell RNA-seq datasets. In general, most reads are located in a region without a feature or within a protein coding gene. However, a high percentage of reads in human bulk RNA-seq datasets are located in other gene types, more specifically mitochondrial genes, due to the high source expression of these genes.

### 3.4 Applying Scavenger recovery on single-cell RNA-seq data

Single cell RNA-sequencing (scRNA-seq) is fast becoming a mainstream method for transcriptomics analysis due its ability to elucidate transcriptional heterogeneity of individual cells. However, there are a number of challenges when dealing with scRNA-seq datasets due to systematically low read counts, as a result of the small amount of transcripts which are captured during library preparation, and a high degree of technical noise [14]. Given Scavenger’s ability in recovering false-negative non-recovered reads in normal bulk RNA-seq datasets and the effect it has on downstream analyses, we hypothesise that recovery of unaligned reads in scRNA-seq datasets with Scavenger will likely have a greater impact on downstream analysis due to limited amount of reads available, while also helping with reducing technical noise. To test this hypothesis, 80 randomly selected samples were collected from a mouse brain scRNA-seq dataset and which are then aligned with STAR, followed by recovery of reads with Scavenger. The scRNA-seq samples have an average read depth of ~2.3 million reads (after pre-processing), with ~57.3% of the reads able to be aligned in the source execution (Table 6). Scavenger was only able to recover 0.6% of the unaligned reads, corresponding to an increase of ~5,400 reads. The low number of reads which are able to be successfully recovered by the Scavenger pipeline is likely due to the low number of aligned in reads in source alignment, which provides less information that Scavenger can utilise during the follow-up execution.

As per the norm for scRNA-seq datasets, the number of genes with nonzero read counts is much lower compared to the number of non-expressed genes in bulk RNA-seq datasets, averaging 5,800. Of these expressed genes, only 12% of the genes (~700) have an increase in read counts, with the majority of these genes having little difference in expression and ~12 genes having a fold-change difference greater than 1 (Figure 8A). Unlike in bulk RNA-seq datasets, genes with substantial difference after recovery range from lowly expressed genes up to highly expressed genes, though genes with the greatest difference in expression are still those with low expression in the source alignment (Figure 4). Furthermore, a different pattern was also observed in the types of genes which have substantial difference in scRNA-seq datasets, with the protein coding category being the majority, followed by the pseudogene category (Figure 8B). The difference in pattern is likely due to comparatively higher abundance of protein coding genes and the low capture efficiency of scRNA-seq methods, meaning that reads from pseudogenes are less likely to be captured and therefore rescued. This can be seen from the much lower number of pseudogenes expressed in scRNA-seq dataset (~150) compared to bulk RNA-seq datasets (~3,500).

**Figure 8:**
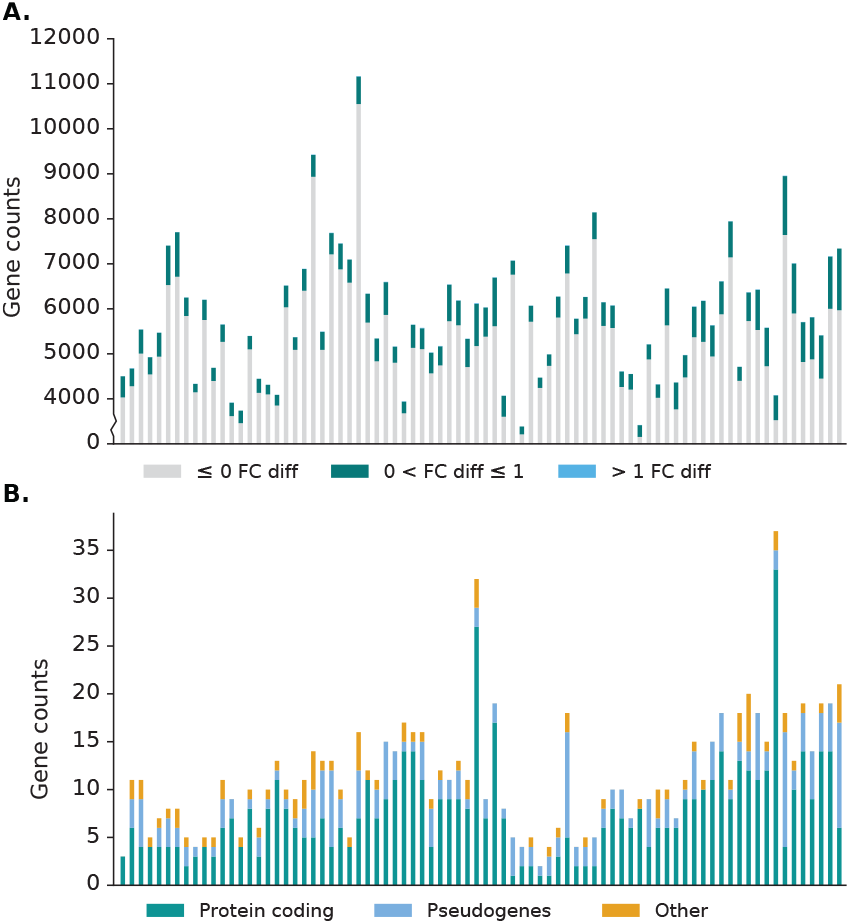
Effect of Scavenger read recovery on gene expression for single-cell RNA-seq datasets. A. The number of genes whose reads are recovered by Scavenger, categorised based on the fold change in normalised expression (RPM) between source alignment and after Scavenger recovery. B. The number of genes with more than 1 fold change in normalised expression categorised based on their gene types.

## 4 Discussion

The false-negative non-alignment problem is a prevalent problem in many of the published RNA-seq alignment tools, resulting in loss of information from incorrectly unaligned reads. To help solve the false-negative nonalignment problem, we have developed Scavenger – a pipeline for recovery of unaligned reads using a novel mechanism based on sequence similarity between unaligned and aligned reads. Scavenger utilises the follow-up execution concept adapted from our previous work on metamorphic testing to find aligned reads from the source execution which have similar sequences to the unaligned reads by aligning the aligned reads against unaligned reads. The location of the aligned reads are then used as a guide to re-align the unaligned reads back to the reference genome using either BLAST or the original alignment tool depending on if the putative location originates from unspliced or spliced alignment, respectively, to ensure that splicing information is retained in recovered reads.

We have applied Scavenger on simulated datasets with varying degrees of complexity and showed that Scavenger is able to recover unaligned reads across all complexity levels with a reasonably high degree of accuracy. In particular, Scavenger is able to recover the most amount of reads in datasets that exhibit a high degree of complexity where read sequence is more divergent compared to the reference genome. We further show that although alignment of reads with optimised parameters are able produce a higher number of aligned reads compared to after recovery with Scavenger, the reads recovered by Scavenger have high degree of overlap to reads recovered with parameter optimisation. The lower number of reads recovered by after Scavenger is a result of Scavenger using information from aligned reads to find putative locations for unaligned reads, meaning that Scavenger is unable to recover reads from region with no alignment – unlike parameter optimisation which does not have the same limitation. Given the non-trivial difficulty of performing parameter optimisation on real datasets, we recommend the use of Scavenger as an alternative to help with recovering incorrectly unaligned reads.

There are a number of possible factors which may contribute to the false-negative non-alignment problem. One such factor is the divergence between the reference genome and the personal genome, leading to higher mismatches during alignment of sequenced reads against the reference genome. In order to validate that divergence of genomic sequences result in incorrectly unaligned reads, we devised an experiment whereby RNA-seq datasets from non-reference mouse strains were aligned against the reference mouse strain. We then analysed the reads which were aligned in the source execution against those recovered by Scavenger and showed that Scavenger is able to significantly recover more reads which have a higher number of reported strain-specific SNPs. This result both confirms that divergence of sequences between the reference genome and the personal genome does affect the false-negative non-alignment problems and that Scavenger is able to recover reads which are incorrectly unaligned due to a higher degree of sequence divergence.

As alignment of reads is only the first step in an RNA-seq data analysis, we also investigated the effect of the false-negative non-alignment problem on downstream analyses, in particular on gene expression analysis. After recovery of reads with Scavenger, we show that ~14 genes have more than 1 fold change in expression compared to the source alignment and that these genes are typically genes with low expression. Interestingly, the majority of genes with >1 expression difference belong to the pseudogenes category, indicating that the expression of pseudogenes are likely to be under-reported due to reads from pseudogenes being incorrectly unaligned by the alignment tool.

Given the ability of Scavenger to recover gene expression in normal (bulk) RNA-seq datasets, we then investigated the ability of Scavenger in recovering reads from scRNA-seq dataset as scRNA-seq datasets have the characteristics of having low reads counts and high degree of technical noise. Scavenger recovery affected the expression of 12% of the expressed genes, with ~12 genes having more than 1 fold change in expression. Unlike the bulk RNA-seq dataset, the genes with >1 change in expression range from lowly expressed genes up to highly expressed genes, with the genes belonging primarily to the protein coding category.

The current version of Scavenger supports STAR as the alignment tool for source execution and re-alignment of spliced reads. However, the user can choose to modify the alignment tool utilised by Scavenger with the alignment tool of their choice. Ideally the tool should satisfy the three properties underlying the read recovery pipeline – deterministic alignment, realignability of mapped reads, and non-realignability of unmapped reads – to ensure that the recovered reads are deterministic. To show the extensibility of Scavenger, we have tested Subread, another RNA-seq alignment tool, as a replacement for STAR within the Scavenger pipeline and demonstrated that Scavenger is still able to recover incorrectly unaligned reads with similar performance to STAR (Table 7 and 8). It should be noted that the recovery performance of Subread is different compared to STAR due to the different algorithm employed by Subread for alignment and, potentially, due to Subread violating the deterministic alignment property.

**Table 7:** Alignment statistics for simulated datasets before and after recovery of reads with Scavenger using default parameters for Subread. The result shown is an average from 3 samples.

| Dataset | Source execution |  |  | Scavenger pipeline |  |  | Unaligned reads recovered | % recovered reads correct | % recovered reads incorrect |
| --- | --- | --- | --- | --- | --- | --- | --- | --- | --- |
|  | Aligned correctly | Aligned incorrectly | Unaligned | Aligned correctly | Aligned incorrectly | Unaligned |  |  |  |
| T1 | 9,305,067 | 74,497 | 620,436 | 9,332,335 | 79,653 | 588,012 | 32,424 | 84.1% | 15.9% |
| T2 | 8,985,799 | 87,576 | 926,625 | 9,107,130 | 92,296 | 800,574 | 126,051 | 96.3% | 3.7% |
| T3 | 4,802,130 | 106,487 | 5,091,384 | 4,984,817 | 108,947 | 4,906,235 | 185,148 | 98.7% | 1.3% |

**Table 8:** Alignment statistics for simulated datasets before and after recovery of reads with Scavenger using optimised parameters for Subread. The result shown is an average from 3 samples.

| Dataset | Source execution |  |  | Scavenger pipeline |  |  | Unaligned reads recovered | % recovered reads correct | % recovered reads incorrect |
| --- | --- | --- | --- | --- | --- | --- | --- | --- | --- |
|  | Aligned correctly | Aligned incorrectly | Unaligned | Aligned correctly | Aligned incorrectly | Unaligned |  |  |  |
| T1 | 9,416,480 | 262,926 | 320,594 | 9,419,057 | 264,906 | 316,037 | 4,557 | 56.5% | 43.5% |
| T2 | 9,283,792 | 397,323 | 318,885 | 9,287,022 | 398,775 | 314,203 | 4,682 | 69.0% | 31.0% |
| T3 | 7,111,603 | 2,251,068 | 637,330 | 7,122,864 | 2,251,625 | 625,512 | 11,818 | 95.3% | 4.7% |

